# Chromatin accessibility associated with aquaculture relevant traits in tilapia

**DOI:** 10.1101/2023.02.17.528929

**Authors:** Tarang K. Mehta, Angela Man, Adam Ciezarek, Keith Ranson, David Penman, Federica Di-Palma, Wilfried Haerty

## Abstract

The Nile tilapia (*Oreochromis niloticus*) accounts for ∼9% of global freshwater finfish production however, extreme cold weather and decreasing freshwater resources has created the need to develop resilient strains. By determining the genetic bases of aquaculture relevant traits, we can genotype and breed desirable traits into farmed strains. We developed and optimised ATAC-seq from *O. niloticus* gill tissues to identify regulatory regions accounting for gene expression associated with gill adaptations. We find that SNPs from 27 tilapia species are enriched in noncoding regions, with 95% of accessible gene promoter regions being SNP-containing. Regulatory variants of TF binding sites are likely driving gene expression differences associated with tilapia gill adaptations, and differentially segregate in freshwater and euryhaline tilapia species. The generation of novel open chromatin data integrated with gene expression and genetic variants revealed candidate genes, genetic relationships, and loci associated with aquaculture relevant traits like salinity and osmotic stress acclimation.

## Introduction

Tilapia cichlid fish of the genus *Oreochromis*, native to Africa and the Middle East, are farmed in over 120 countries/territories ^1^. The Nile tilapia, *Oreochromis niloticus*, accounted for 9% of the 49 million tonnes of freshwater finfish produced globally in 2020, the third most of any species ^1^. The exponential growth in tilapia aquaculture production is largely due to their suitability for aquaculture systems and unlike most other finfish species, tilapia can grow and reproduce in many culture systems. However, climate change is leading to extreme weather events, and decreasing freshwater resources. There is a pressing need to develop aquaculture systems based on strains resilient to saline waters and temperature changes. A way forward is to characterise the genetic bases responsible for such adaptive traits enabling selective breeding into farmed strains.

Such an approach involves resequencing populations to map and characterise variation, and identify signatures of selection in genomic regions associated with adaptive differences ^2^. Previous population studies using whole genome resequencing of wild tilapia individuals (*O. niloticus, O. mossambicus*, and red tilapia) and commercial strains reported between 1.3 and 1.43 million single nucleotide polymorphisms (SNPs) ^3,4^, the majority of which are located within noncoding regions of the genome. Most SNPs under selection were found within those noncoding regions (45% intronic and 19% intergenic) and associated with several aquaculture relevant traits like growth, reproduction, and immunity ^3,4^. Only 16 non-synonymous SNPs were identified in the three wild tilapia populations, suggesting selection on functional noncoding regions has played a prominent role during domestication/breeding ^3^.

We recently generated genome-wide sequencing data across 575 individuals of 27 tilapia species from across East Africa, to identify 69 million single nucleotide polymorphisms (SNPs) in the *O. niloticus* genome assessing phylogenomic patterns of recent and ancient hybridisation in tilapia (Ciezarek, A. *et al*., bioRxiv TBC, 2023). Whilst selection in noncoding regulatory regions, including transcription factor binding sites (TFBSs) has been suggested as a key molecular mechanism contributing to the evolutionary diversification of East African cichlids, including Nile tilapia ^5-7^, no study has evaluated the *cis*-regulatory impact of noncoding variation towards adaptive traits in a tilapia phylogeny.

Several studies have provided evidence on the role of *cis*-regulatory elements (CREs) towards morphological diversity ^8-10^, including in mammals ^11^ and fish ^12^. Whilst such studies identified CREs at particular loci, epigenetic sequencing methods can capture chromatin accessibility and TF-binding at the genome-wide level ^13^. Only a handful of studies, all using DNA methylation, have applied epigenetic approaches to study adaptive traits in tilapia ^14^; this includes the regulation of tilapia growth ^15-17^, sexual dimorphism ^18^, and sex determination ^19,20^. ChiP-seq approaches have also been used to map active promoter elements associated with Nile tilapia fin development ^21^. However, such techniques are only able to identify repressive or active marks at specific loci, and in the case of ChiP-seq, direct interactions between a specific protein and DNA. In contrast, the assay for transposase-accessible chromatin using sequencing (ATAC-seq) approach can robustly identify genome-wide chromatin accessibility and TF-occupancy using very few ^22,23^ or even single cells ^24,25^. Despite the robustness of ATAC-Seq and its potential to provide a better annotation of regulatory regions, and an accurate identification of TF-occupancy including binding sites for regulators in Nile tilapia, no such study has been performed to date on tilapia tissues and/or cells.

Here, we aim to identify genes and functional non-coding regions in Nile tilapia by characterising the open chromatin and gene expression landscape of Nile tilapia gill tissue to classify functional noncoding variation associated with environmental tolerance. For this, we characterised the open chromatin landscape of Nile tilapia gill tissue using ATAC-seq, and then identified accessible gene promoter regions with TF footprints that could account for target gene expression in the gill. We integrate this data with noncoding regulatory SNPs from a 27 species tilapia phylogeny (Ciezarek, A. *et al*., bioRxiv TBC, 2023), and identify candidate genes with regulatory variation that could be associated with aquaculture-relevant traits, like salinity tolerance.

## Results

### Accessible gene promoter regions in gill tissue are functionally associated with aquaculture relevant traits

To characterise functional noncoding regions based on chromatin accessibility, we performed high-depth ATAC-seq of gill tissue from three replicate male Nile tilapia individuals, identifying 301,293 open chromatin (accessible) reproducible peaks between replicates (see ‘*Materials and Methods*’), that could be associated to 25,092 (average of 12 accessible peaks per gene) out of 29,552 (85%) annotated Nile tilapia genes ^26^. We first annotated (Supplementary Table S1) and tested the enrichment of accessible peaks in coding and noncoding regions, finding that peaks are mostly enriched (fold enrichment >2.1, adjusted *P-*value <0.05) in both sets of conserved noncoding elements (CNEs) (Fig. 1) that we annotated in the Nile tilapia (*O. niloticus* UMD_NMBU ^26^) reference genome (see ‘*Materials and Methods*’).

**Fig. 1.**
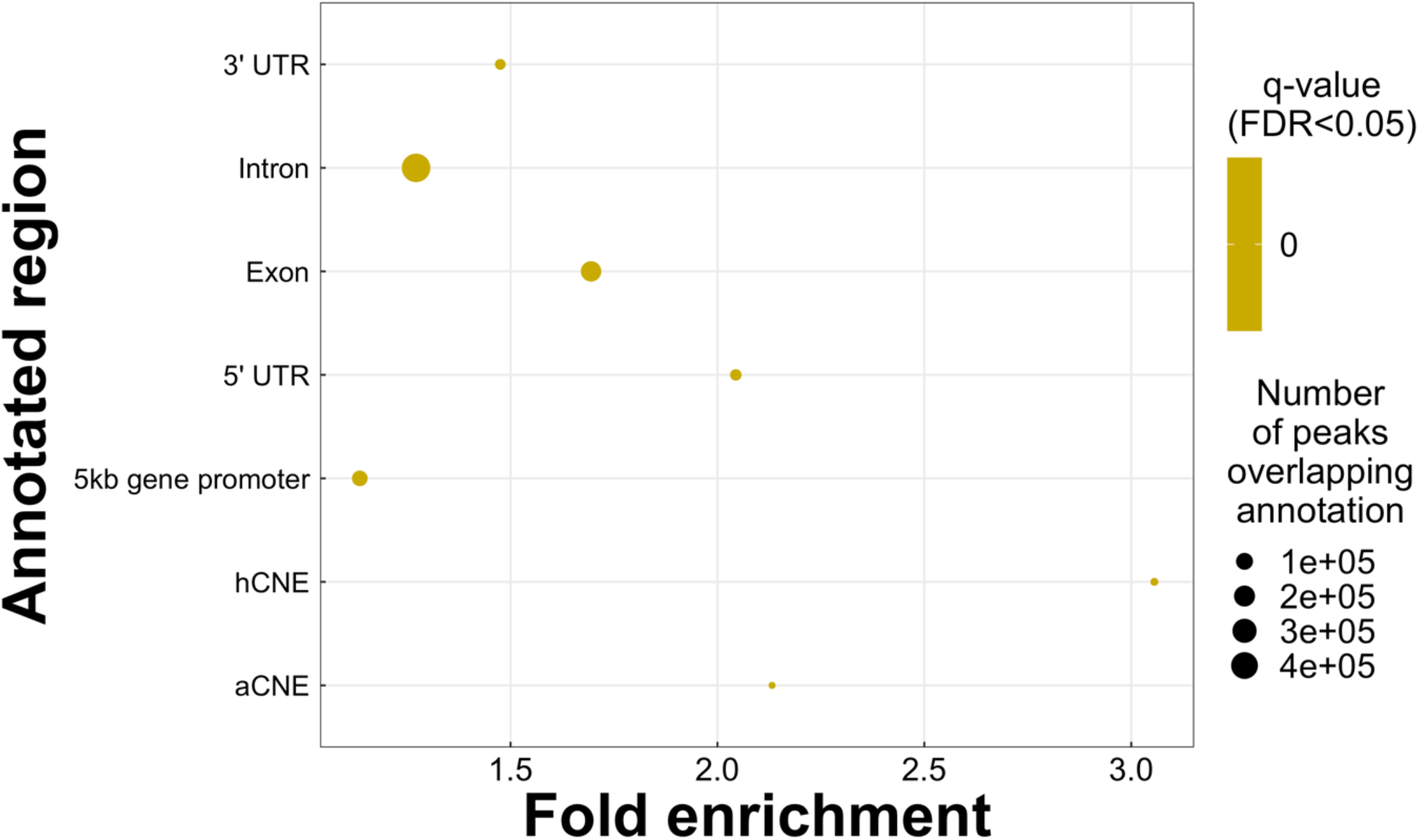
Annotation of open chromatin peaks in Nile tilapia gill. Fold enrichment of open chromatin (accessible) peaks overlapping coding and noncoding regions in the Nile tilapia genome. Circles show enriched annotated region (y-axis) of significance (FDR <0.05, to right) and fold enrichment (x-axis) values of all annotated peaks. Number of peaks overlapping each annotation shown by size of each circle.

We focused on gene promoter regions, taken as up to 5 kb from the transcription start site (TSS) ^7^, as they harbour regulatory binding sites e.g., TFBSs that have functionally diverged ^6,7^, and can be associated with gene expression differences in Nile tilapia and East African cichlids ^7^. Out of the 301,293 open chromatin (accessible) peaks, 127,602 accessible peaks (average size of 464 bp) were found within gene promoters of 20,248 genes in gill tissue, with missing genes being better associated to lymphoid tissues (see ‘*Supplementary Information’*, Supplementary Fig. S1 – S2).

### Accessible gene promoter regions could account for expression changes of genes associated with osmoregulation and salinity tolerance

To examine any correlation between gene transcription and accessible gene promoter regions, RNA-seq data was generated from the same gill tissue sample of the three replicates and gene expression measured using transcripts per million (TPM) (see ‘*Materials and Methods*’). Of the 29,552 Nile tilapia genes in the genome, we identified and calculated the expression (as TPM) of 20,717 (70%) genes. We found 96,784 (76%) accessible peaks could be associated to the expression of 14,842 genes.

We first tested whether these peaks are better correlated to their corresponding gene expression based on proximity to the transcription start site (TSS) (Fig. 2a). For this, accessible peaks are categorised into four intervals of either 500 bp (12,978 peaks), 1 kb (27,975 peaks), 2kb (48,108 peaks) or up to 5 kb (127,602 peaks) from the genes TSS. We found similar weak positive correlation (R = 0.13 to 0.14, Spearman *p* < 2.2e-16) for all four intervals (TSS peak up to 500 bp, 1 kb, 2 kb and 5 kb) (Fig. 2a) and thus, we focus on all (159,708) accessible gene promoter peaks (up to 5 kb from the TSS). A higher correlation might not be observed in our datasets as accessible regions are not always associated with gene activity and can instead, be associated with repressed or poised for activation genes ^27-30^.

**Fig. 2.**
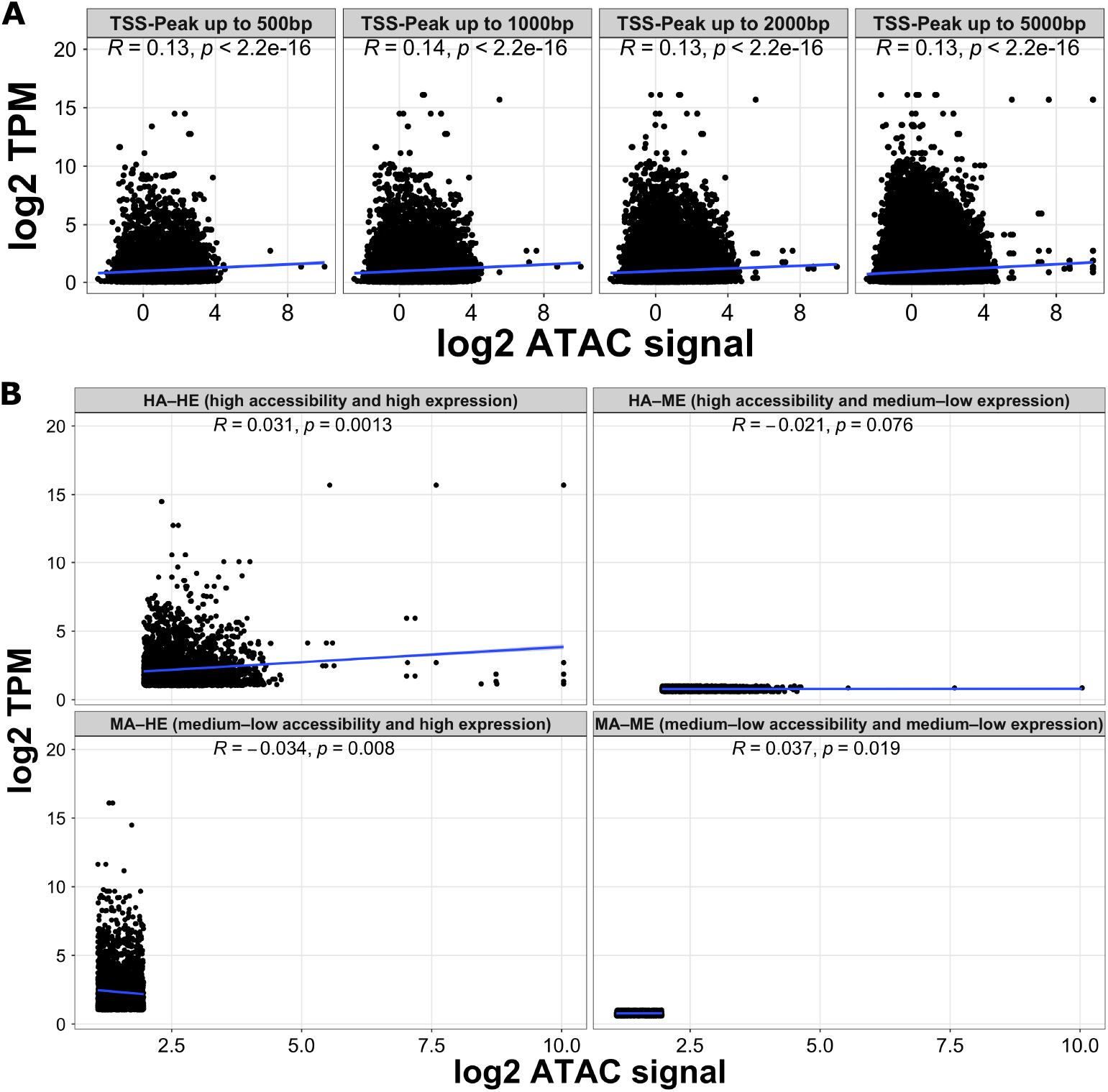
Correlation of accessible peaks accounting for gene expression change in Nile tilapia gill. **(A)** Average *log2* gene expression (as transcripts per million - TPM, y-axis) and average *log2* ATAC signal (as gene promoter peak counts, x-axis) for each gene across the three replicates, split into four categories based on peak summit distance to transcription start site (TSS). Correlation co-efficient (*R*, blue trendline) and *p-value* shown for each category. **(B)** Average *log2* gene expression (as transcripts per million - TPM, y-axis) and average *log2* ATAC signal (as all gene promoter peak counts i.e., TSS-peak up to 5000 bp category in Fig. 2a, x-axis) for each gene across the three replicates, split by high accessibility and high expression to left (HA-HE - both average TPM and ATAC signal values are above the 70^th^ percentiles) and to right, medium-low accessibility and high expression (MA-ME - average ATAC signal count less than the 50^th^ percentile and TPM higher than the 70^th^ percentile). Correlation co-efficient (*R*, blue trendline) and *p-value* shown for each category.

Based on an approach devised previously ^30^, we explore the relationship of accessible peaks and gene expression by assessing genes with combinations of high accessibility (HA) or medium–low accessibility (MA), and either highly expressed (HE) or medium–low expression (ME) (see ‘*Materials and Methods*’). After categorising gene peaks and expression, we were left with a total of 27,536 (22% of 127,602) accessible peaks associated to the expression of 4,509 unique genes. These could be placed into the four categories, with most genes (1,650) and peaks (10,488) in the HA-HE category, and the remaining being either HA-ME (6,961 peaks and 1,081 unique genes), MA-HE (5,990 peaks and 1,770 unique genes), or MA-ME (4,097 peaks and 1,164 unique genes). Whilst the HA-ME genes exhibit no correlation (R = -0.021, Spearman *p* = 0.076), and the MA-HE genes exhibit significant weak negative correlation (R = -0.034, Spearman *p* = 0.008), both the MA-ME (R = 0.037, Spearman *p* = 0.019) and HA-HE genes (R = 0.031, Spearman *p* = 0.0013) exhibit significant weak positive correlation (Fig. 2b). Among the 1,650 HA-HE genes, we identify genes associated with aquaculture-relevant traits, including those enriched (FDR < 0.05) for the GO terms ‘cellular process’ e.g., *prlr1* involved in tilapia salinity tolerance ^31^ and *clcn2* an osmoregulatory gene involved in adaptation to saline-alkali challenge in tilapia ^32^, and ‘cellular respiration’ e.g., *cox4* involved in aerobic metabolism changes under salinity challenge in tilapia gills ^33^ (Supplementary Fig. S3).

### Nile tilapia SNPs are enriched in noncoding regions

Since we previously found that discrete variation in regulatory regions disrupt regulatory network interactions for adaptive trait genes in Nile tilapia ^6,7^, we tested whether functional noncoding SNPs could be associated with the regulatory activity of genes involved with aquaculture-relevant traits, like environmental tolerance. Building upon our previous work of identifying 69 million (69,064,774) SNPs across 27 tilapia species (Fig. 3a, Supplementary Table S2) mapped to the Nile tilapia genome (Ciezarek, A. *et al*., bioRxiv TBC, 2023), we identified that these SNPs are significantly enriched in noncoding than coding (exonic) regions (Fig. 3b). Compared to exonic regions (fold enrichment of 0.9, adjusted *P-*value <0.05), the SNPs are most enriched in either intronic or up to 5 kb gene promoter (fold enrichment of 1.1, adjusted *P-*value <0.05) regions (Fig. 3b).

**Fig. 3.**
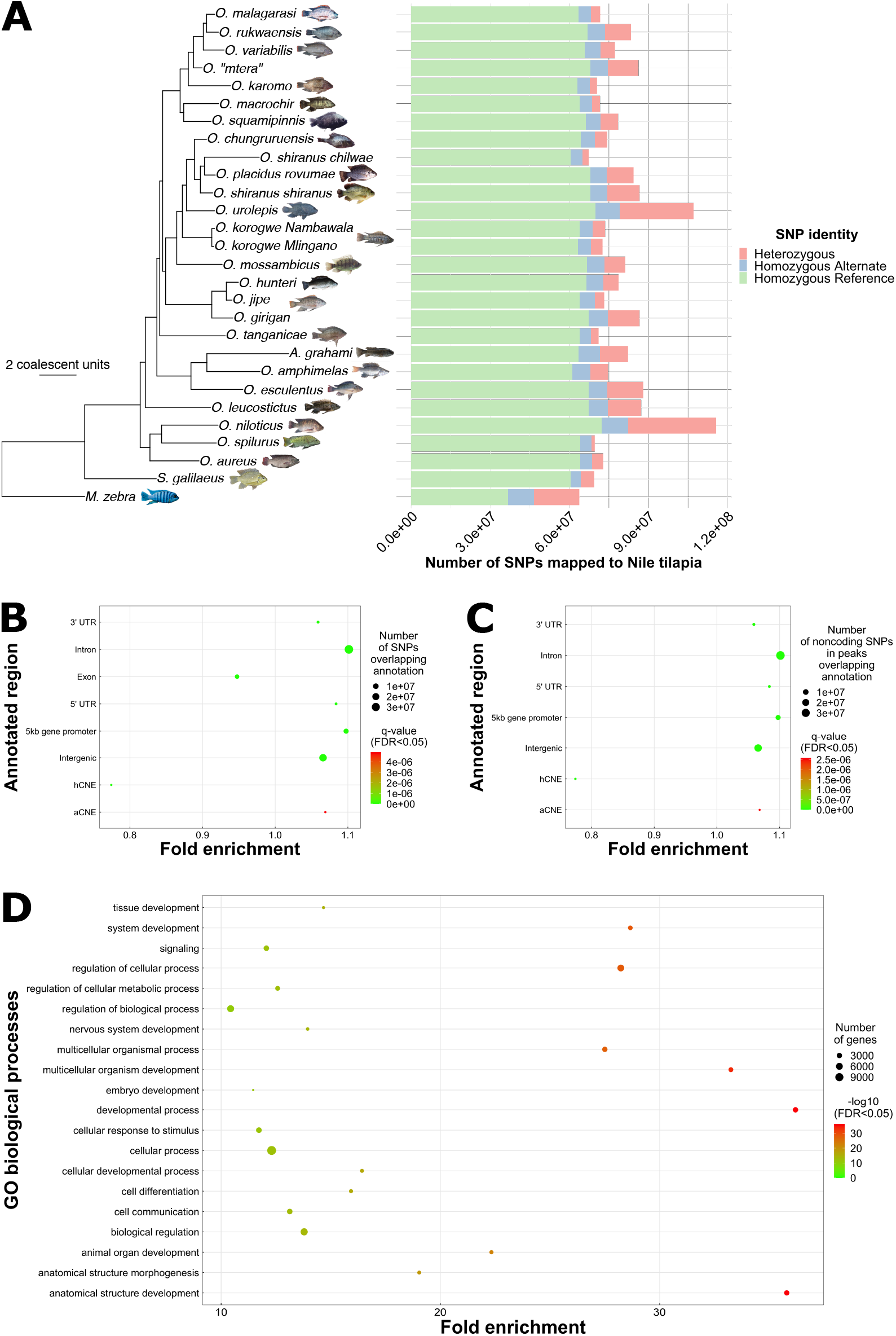
Single nucleotide polymorphisms (SNPs) in the Nile tilapia genome. **(A)** Coalescent phylogeny of 27 tilapia species and 2 outgroups inferred using ASTRAL (Ciezarek, A. *et al*., bioRxiv TBC, 2023) with number and identity of SNPs in each species based on mapping to the Nile tilapia genome. Phylogenies inferred based on 14,535 10 kb windows across the genome. Units = coalescent units; All posterior probabilities = 1.0. Species photo credits: George Turner, Martin Genner, Antonia Ford, Benjamin Ngatunga, Geraldine Kavembe, and Fishbase (Luc de Vos, Brain Gatwicke, Andrew Nightingale, and Royal Museum for Central Africa). **(B)** Fold enrichment of SNPs overlapping coding and noncoding regions in the Nile tilapia genome. Circles show enriched annotated region (y-axis) of significance (FDR<0.05, heatmap to right) and fold enrichment (x-axis) values of all SNPs mapped to the Nile tilapia genome. Number of SNPs overlapping each annotation shown by size of each circle. **(C)** Fold enrichment of SNPs overlapping open chromatin (accessible) peaks in noncoding regions of the Nile tilapia genome. Circles show enriched annotated region (y-axis) of significance (FDR <0.05, heatmap to right) and fold enrichment (x-axis) values of all SNPs in annotated noncoding peaks. Number of noncoding SNPs in annotated noncoding peaks shown by size of each circle. **(D)** Gene ontology (GO) biological process enrichment of accessible gene promoter regions containing SNPs in the Nile tilapia genome. GO terms are subset for enrichment of -log10 (FDR < 0.05) of more than 10, with the full figure provided as Supplementary Fig. S5. Circles show enriched term (y-axis) of significance (FDR <0.05, heatmap to right) and fold enrichment (x-axis) values of GO terms. Number of genes with promoter regions containing SNPs shown by size of each circle.

To characterise functional noncoding variation, we find that of the 5,470,066 SNPs in Nile tilapia gene promoter regions, 1,083,195 (20%) SNPs overlapping 121,404 of the 127,602 (95%) accessible peaks are significantly enriched (fold enrichment of 1.1, adjusted *P-*value <0.05) (Fig. 3c). Of the 20,248 accessible gene promoters, 20,101 (99%) have SNPs in accessible regions, representing 68% of all annotated Nile tilapia genes ^26^. Most SNPs are rare variants (Minor Allele Frequency, MAF <= 0.1) with 1% more rare variants found in noncoding regions that are accessible than not (Supplementary Fig. S4, see ‘*Supplementary Information’*). Using the 127,602 accessible peaks in gene promoter regions, we find that the topmost (-log10 FDR < 0.05 of >=10) enriched GO biological processes of 20,248 accessible gene promoter regions harbouring SNPs could be functionally relevant to aquaculture traits like, for example, signalling, cell communication, and cellular response to stimulus (Fig. 3d, Supplementary Fig. S5). In summary, nearly three-quarters of all Nile tilapia genes have putative functional noncoding variation at gene promoter regions that could be associated with the regulation of aquaculture traits relevant to gill function like, for example, a response to stimulus.

### Discrete variation in accessible TFBSs is likely driving gene expression associated with tilapia gill adaptations

We hypothesised that genetic variation between Nile tilapia and the 26 other tilapia species (Fig. 3a) could be used to identify functional noncoding variation based associated with aquaculture traits that differ between the species. For example, genetic variation in gene promoter TFBSs between Nile tilapia (a freshwater species) and any euryhaline species e.g., *A. grahami, O. mossambicus* or *O. urolepis* with exclusive homozygous or heterozygous alternate sites could indicate positive selection acting on discrete mutations, that can be functional based on chromatin accessibility in Nile tilapia gill tissue, and subsequently associated to target gene expression of gill-specific traits relevant to aquaculture e.g., salinity tolerance. To prioritise predicted TFBSs (see ‘*Materials and Methods*’ and ‘*Supplementary Information’*), we use the best described method for ranking predictions ^34^ by overlapping predicted TFBSs with the number of reads (tag count, TC) in the footprint, and then rank TFBSs using the bit-score of the motif match within ranked tag counts. Based on a mean bit-score of 11.5 and mean tag count of 111.9, we identify 6,195,938 TFBSs above the means (Fig. 4a). In gene promoter regions, we identify 246,296 TF footprints with 2,491,107 non-redundant TFBSs (see ‘*Materials and Methods*’). Based on a mean bit-score of 11 and mean tag count of 152.6, we identify 398,995 (16%) TFBSs above the means in gene promoter regions.

**Fig. 4.**
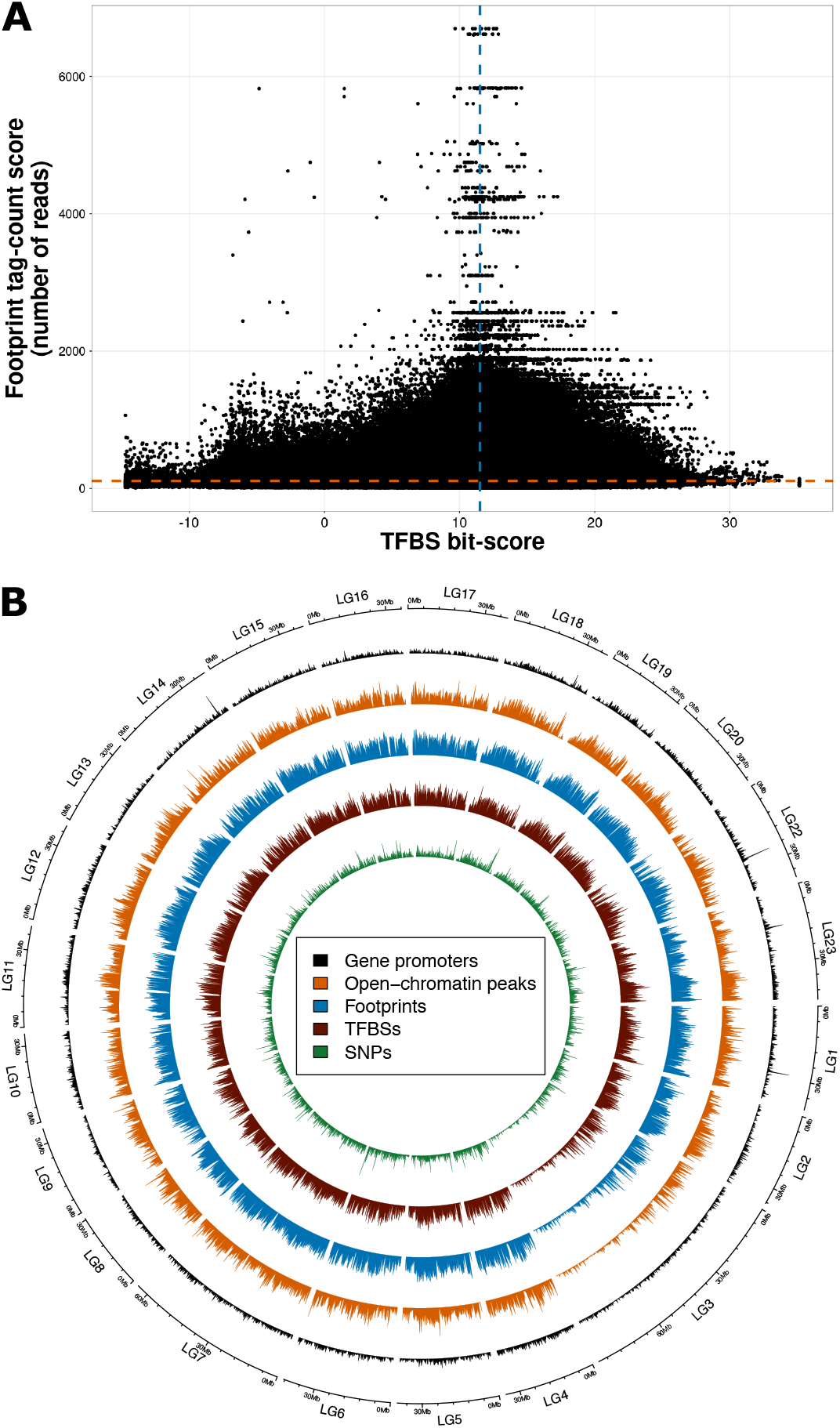
Prioritising genetic variation of TFBSs in accessible peaks accounting for gene expression change in Nile tilapia gill. **(A)** Tag-count (TC) score of TF footprints (y-axis) and TFBS bit-score of motif match (x-axis) of genome-wide predicted TFBSs based on footprints in accessible gill peaks. **(B)** Circos plot of open-chromatin peaks (orange), footprints (blue), TFBSs (maroon), SNPs (green), and gene promoter regions (black) on each Nile tilapia chromosome (outer track).

Using TFBSs in accessible gene promoter regions, we identified a total of 3,398,768 transcription factor (TF) – target gene (TG) relationships in Nile tilapia. We then identify discrete mutations in TFBSs to characterise conservation or divergence and therefore, whether the 3,398,768 Nile tilapia TF-TG relationships could be present or absent in the other 26 species. First, we map open-chromatin peaks, footprints, TFBSs and SNPs to gene promoter regions on each Nile tilapia chromosome (Fig. 4b). In total, we identify 7,029,307 (11% of total 69,064,774) SNPs overlapping accessible gene promoter regions, TF footprints, and TFBSs with all (2042) TF motifs having at least one discrete mutation in 18,899 genes (64% of the 29,552 Nile tilapia genes in the genome ^26^). More than half (62% - 11,790) of the 18,899 genes have SNPs in ‘prioritised’ TFBSs where the bit-score of motif match and tag count are above the means of 11 and 152.6 respectively. Of the 18,899 genes, 1,562 (8%) have high accessibility and high expression (HA-HE), which makes up the majority (95%) of all 1,650 HA-HE genes. This indicates that discrete variation in accessible gene promoter TFBSs could be a core factor driving high levels of gene expression associated with environmental tolerances of the gill. Using both measures, we narrowed this down to 1,168 genes (71% of all HE-HE genes or 6% of all genes with a SNP in an accessible gene promoter TFBS) that are categorised as 1) having a SNP in a ‘prioritised TFBSs’ where the bit-score of motif match and tag count are above the mean of all gene promoter TFBSs; and 2) being a ‘HA-HE gene’ based on peak-expression correlations. To further narrow down the 1,168 genes, we focus on genes 3) with ‘genetic variation’ that exists between species exhibiting a gradient of environmental tolerance e.g., freshwater versus saline water; and 4) that are enriched for biological process GO terms that are associated with gill adaptations of traits with aquaculture relevance e.g., response to stimulus (Fig. 3d, Supplementary Fig. S5) or cellular process (Supplementary Fig. S3). We use these four described criteria for identifying examples of candidate genes that are likely involved in gill function associated with environmental tolerance in tilapia.

### Discrete mutations at accessible regulatory sites of adaptive trait genes are associated with environmental tolerance networks in tilapia species

Using the four described criteria (prioritised TFBSs, HA-HE gene, genetic variation between diverse species, and aquaculture relevant GO terms) on the set of 1,168 genes, we first identified STAT1-*prlr1* (Fig. 5) as a candidate TF-TG relationship that could be involved in gill adaptations associated with tilapia environmental tolerance. The STAT1 TFBS is proximal (<400 bp) to the *prlr1* TSS (Fig. 5a-b) and with a significant signal of activity (Fig. 5c), has a few standard deviations higher bit-score of motif match (11.04 ± 2.61 SD) and tag count (152.6 ± 0.12 SD) than the means of all predicted gene promoter TFBSs. Differences in the regulation of prolactin (PRL) and its receptors (PRLRs) have been shown to underly variation in salinity tolerance of tilapias ^31^, and are differentially expressed between *A. grahami* and the freshwater species, *O. leucostictus* gill transcriptomes ^35^. For this reason, genetic variation at the TFBS for STAT1 could account for *prlr1* gene expression changes associated with salinity response in different tilapia species. We identified a rare variant (LG7 position 18107947; MAF=0.03) at the third position in the predicted STAT1 TFBS (Fig. 5b and Fig. 5d). The rare variant is homozygous alternate G/G (n=14) and heterozygous alternate T/G (n=9) in the highly adapted extremophile species, *A. grahami*, but homozygous reference T/T in Nile tilapia and the other 25 tilapia species (Fig. 5d). STAT1 is a transcriptional activator that can be phosphorylated via osmotic stress ^36^; this suggests that STAT1 could be a key regulator of *prlr1* gene expression in most tilapia species gills in response to osmotic stress however, the loss of STAT1 regulation of *prlr1* could necessitate gill adaptations to extreme alkaline and saline waters that are natural to *A. grahami* ^37,38^.

**Fig. 5.**
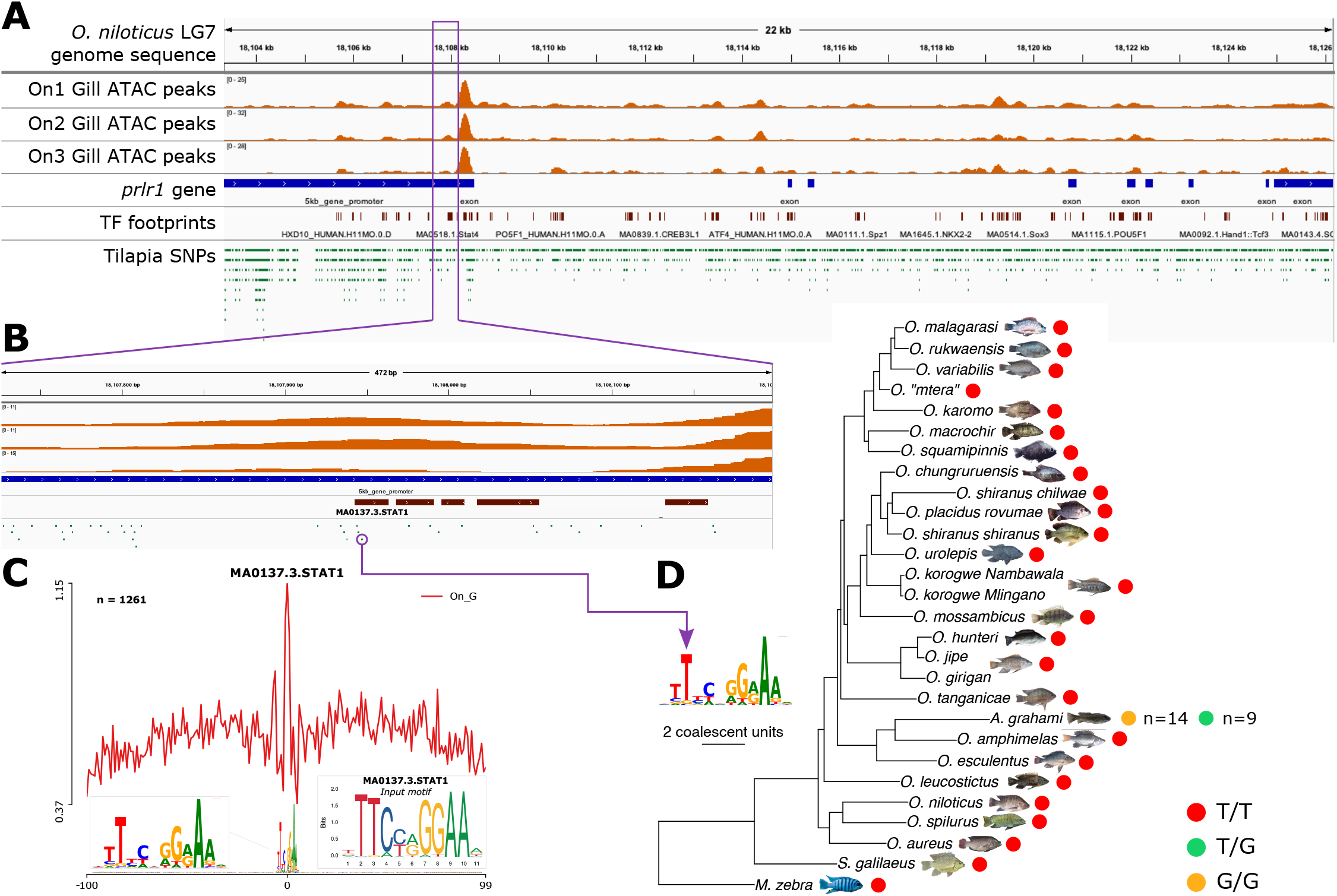
Accessible peaks in the Nile tilapia *prlr1* gene promoter with variation at the STAT1 TF footprint. **(A)** Nile tilapia track of the *prlr1* gene sequence (blue annotations) in LG7 with accessible peaks across three biological gill replicates (orange peaks), TF footprints (maroon marks) and tilapia SNPs (green marks). **(B)** Expanded *prlr1* gene promoter region (blue annotation) of accessible peaks (in orange) across three gill replicates above the STAT1 TF footprint (maroon mark) and containing a SNP (green mark with purple circle). **(C)** Line plot profile of the STAT1 TF footprint (n=1261) across Nile tilapia gene promoter regions showing positional (x-axis) signal of activity (red line, y-axis) centred on the footprint (at position 0) with flanking sequence. Expanded STAT1 TF footprint shown to left and original input motif used for predicting the site shown to right. **(D)** Variation from the SNP in Fig. 5b demarcated in the STAT1 TF footprint with homozygous reference (red dots), homozygous alternate (green dots) and heterozygous (yellow dots) alleles for the SNP shown for each species in the tilapia phylogeny (based on Fig. 3a).

In another example, we identify variation in the gene promoter region of *cldnh* (also referred to as *cldn3-like* and *cldn28a*), associated with expression differences in stickleback ecotypes ^39^ and euryhaline tilapia ^40^ under salinity challenge, as well as involved in permeability changes associated with salinity acclimation in freshwater tilapia species too ^41^. Specifically, we identify SREBF2-*cldnh* (Fig. 6) as another candidate TF-TG relationship that could be involved in gill adaptations associated with salinity acclimation, especially of freshwater species. The SREBF2 TFBS is proximal (<330 bp) to the *cldnh* TSS (Fig. 6a-b) with a significant signal of activity (Fig. 6c), and a few standard deviations higher bit-score of motif match (11.04 ± 0.4 SD) and tag count (152.6 ± 0.3 SD) than the means of all predicted gene promoter TFBSs. We identified a rare variant (LG10 position 17991869; MAF=0.07) at the 4th position in the predicted SREBF2 TFBS (Fig. 6b and Fig. 6d). We identify the rare variant is mostly homozygous alternate G/G (n=19) and heterozygous alternate A/G (n=14) in the freshwater species, *O. esculentus*, but homozygous reference A/A in Nile tilapia, *O. esculentus* (n=4) and most individuals, except *O. “mtera”*, of the other 25 tilapia species (Fig. 6d). Despite being a freshwater species, *O. esculentus* individuals with heterozygous and homozygous alternate genotypes were collected from saline/alkaline water bodies, namely Hombolo Dam ^42^, Lake Malimbe ^43^, and Lake Rukwa ^44,45^. Such genetic variation at the SREBF2 TFBS could therefore account for differential expression of the *cldnh* gene, which has been previously shown to reorganise gill tight junctions in response to salinity acclimation of euryhaline ^40^ and freshwater ^41^ tilapia. Discrete mutations in regulatory sites of aquaculture relevant genes could therefore be driving gene regulatory evolution associated with salinity acclimation of the gills.

**Fig. 6.**
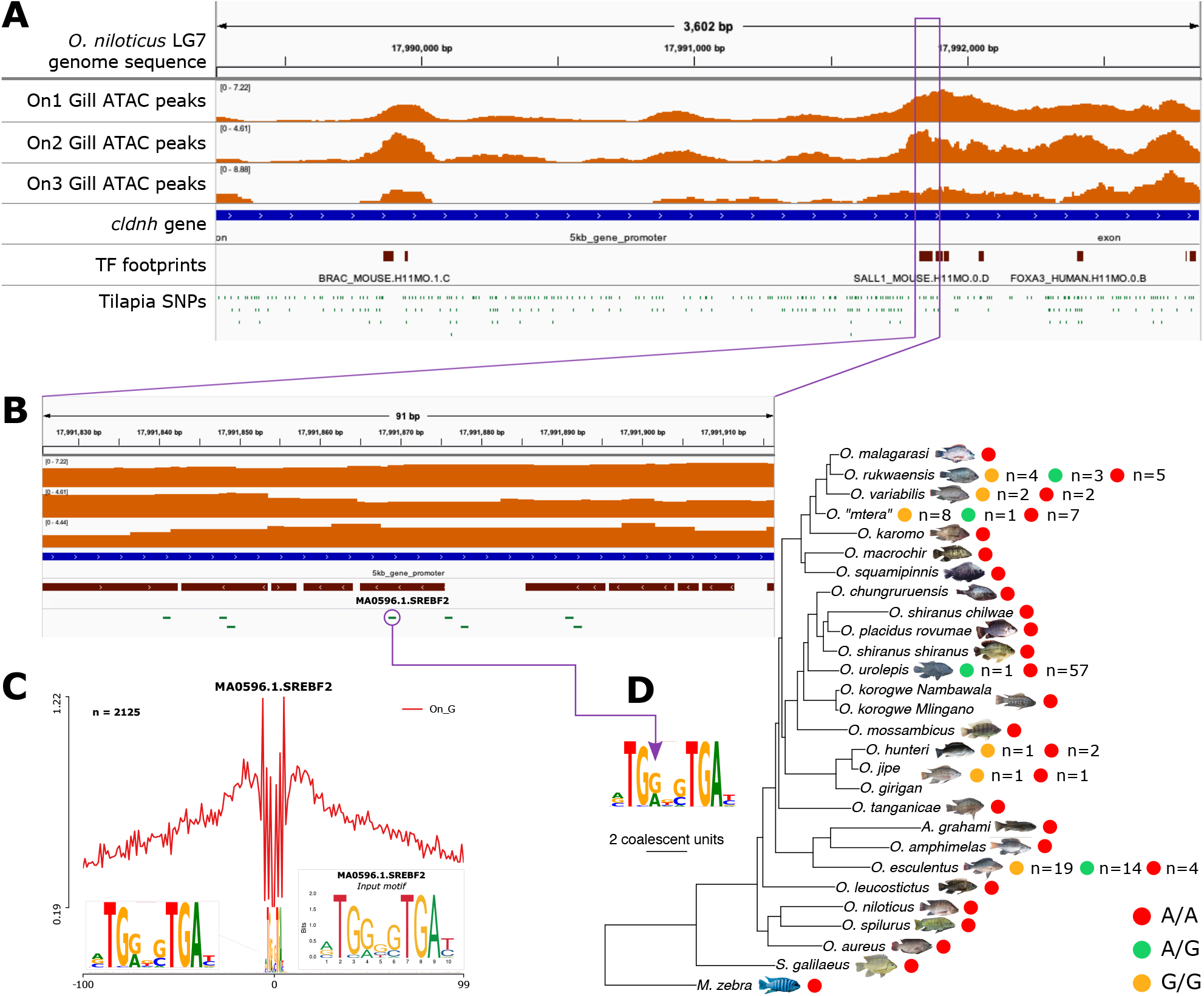
Accessible peaks in the Nile tilapia *cldnh* gene promoter with variation at the SREBF2 TF footprint. **(A)** Nile tilapia track of the *cldnh* gene sequence (blue annotations) in LG10 with accessible peaks across three biological gill replicates (orange peaks), TF footprints (maroon marks) and tilapia SNPs (green marks). **(B)** Expanded *cldnh* gene promoter region (blue annotation) of accessible peaks (in orange) across three gill replicates above the SREBF2 TF footprint (maroon mark) and containing a SNP (green mark with purple circle). **(C)** Line plot profile of the SREBF2 TF footprint (n=2125) across Nile tilapia gene promoter regions showing positional (x-axis) signal of activity (red line, y-axis) centred on the footprint (at position 0) with flanking sequence. Expanded SREBF2 TF footprint shown to left and original input motif used for predicting the site shown to right. **(D)** Variation from the SNP in Fig. 6b demarcated in the SREBF2 TF footprint with homozygous reference (red dots), homozygous alternate (green dots) and heterozygous (yellow dots) alleles for the SNP shown for each species in the tilapia phylogeny (based on Fig. 3a).

## Discussion

Most (∼90%) tilapia aquaculture production is based on Nile tilapia ^46^ owing to its high growth and reproduction rate in several culture systems. However, with climate change leading to extreme weather and human competition decreasing freshwater resources, there is a crucial need to breed tilapia strains that are resilient to broad environmental conditions. One method to determine the genetic bases responsible for environmental traits e.g., salinity tolerance, involves firstly characterising genetic variation and signatures of selection associated with adaptive traits in resequenced populations ^2^ and then secondly, using epigenetic sequencing e.g., ATAC-Seq to annotate functional genomic regions and RNA sequencing to characterise whether these loci can drive gene expression change. Despite the impact of noncoding regulatory evolution towards the diversification of cichlids, including Nile tilapia ^5-7^ and breeding wild tilapia populations ^3,4^, no previous work has applied ATAC-Seq to accurately identify transcriptional regulators in tilapia tissues and/or cells, let alone classify functional noncoding variation associated with adaptive traits.

By performing high-depth ATAC-seq of Nile tilapia gill tissues, we identify 301,293 accessible (open chromatin) regions in three replicates that can be associated to 85% of all annotated Nile tilapia genes ^26^. No previous work has carried out ATAC-seq in gill tissue however, we identify more peaks in our study compared to 11 other zebrafish tissues (66k-180k peaks) but fewer than the 436k merged non-redundant peaks across all tissues ^47^. The identification of more single tissue peaks than zebrafish is possibly due to tissue-specific differences and sampling of more replicates. More than 40% (127,602) of the accessible regions can be found within gene promoter regions of 20,248 genes, with genes without a reproducible peak being better associated to functions of lymphoid tissues not sampled here.

We were able to associate 96,784 (76%) accessible gene promoter regions to the expression of 14,842 genes. After determining no difference in weak positive correlation of accessible peaks to gene expression based on proximity to the TSS, we found that most genes (1,650 out of 4,509) could be categorised as highly accessible and highly expressed (HA-HE) based on signals and counts above the 70th percentiles. The HA-HE genes are enriched for genes involved in salinity acclimation however, the overall set exhibits weak positive correlation of accessibility and gene expression. A higher correlation might not be observed in our datasets as accessible regions are not always associated with gene activity and can instead, be associated with repressed or poised for activation genes ^27-30^. Like previous work in *C. elegans* ^*27*^, this would need to confirmed using relevant (H3K27me3 and H3K4 methylation) ChiP-Seq marks.

Whilst our previous work focused on regulatory variation in a phylogeny of five cichlid species, including Nile tilapia ^6,7^, no previous work has examined functional noncoding variation in the tilapia phylogeny. Building upon our recent work of identifying 69 million SNPs in the Nile tilapia genome based on resequencing populations of 27 tilapia species (Ciezarek, A. *et al*., bioRxiv TBC, 2023), we found SNPs are enriched in noncoding regions and more specifically, gene promoter regions. We identified that gene promoter regions with accessible SNPs are enriched for processes associated with gill function e.g., response to stimulus, and therefore conclude that much like the adaptive radiations of East African cichlids ^6,7^, discrete variation in regulatory regions can also drive gene regulatory rewiring of tilapia adaptive trait genes. This is supported by the fact that nearly all (99%) accessible gene promoter regions have a SNP with most (90%) being a rare variant. Gene promoter regions are enriched for TFBSs ^7^, as are accessible regions ^47,48^, and thus, variation, including rare variation ^49^, at TFBSs is a major target of genetic selection ^50^ and a key contributor to adaptive phenotypes ^6,7,51^. Using TFBSs in accessible gene promoter regions, we identified a total of 3,398,768 transcription factor (TF) – target gene (TG) relationships in Nile tilapia. As we use experimental data, this is less than the 5,900,174 TF-TG edges we *in silico* predicted in Nile tilapia in our previous work ^7^, but similar to the 3,505,491 TF-TG relationships found in human based on *in silico* predictions overlapping ENCODE footprints ^52^.

Using a stringent criterion to identify candidate TF-TG relationships, involving prioritising statistically significant TFBSs with genetic variation between ecologically divergent species and genes that exhibit high accessibility and high expression compared to all genes, we identified 1,168 genes that could be associated to aquaculture relevant traits, like salinity acclimation. Some tilapia species can thrive in adverse environmental conditions, including varying levels of salinity and alkaline water. Nile tilapia is intolerant to high salinity (optimal is up to 16 parts per thousand, ppt) ^53^ and requires freshwater (pH 6-9) ^54,55^ whereas euryhaline species like *Alcolapia grahami*, nested within the *Oreochromis* genus ^56^, have adapted to extreme environments of pH 9-11.5 and high salt concentrations of >20 ppt ^37,38^. In *prlr1*, a gene associated with tilapia salinity response ^31^, a rare genetic variant between Nile tilapia and *A. grahami* in the STAT1 TFBSs could account for osmotic stress response in most tilapia species, but the loss of this site could enable gill adaptation in extremophiles like *A. grahami*. This is further supported by the role of STAT1 as an osmotic stress induced transcriptional activator ^36^ and differential expression of prolactin receptors in gills of *A. grahami* and freshwater tilapia species ^35^. On the other hand, the introduction and acclimation of freshwater species, like *O. esculentus*, to saline/alkaline waters e.g., Lake Rukwa in Tanzania ^44,45^, could be driven by genetic variation at the SREBF2 TFBS, and account for differential expression of *cldnh*, a gene responsible for reorganising gill tight junctions in response to salinity exposure of freshwater tilapia ^41^. Salinity acclimation first requires the detection of osmotic changes and then secondly, a physiological response to restore osmotic homeostasis. Accordingly, excessive ions are secreted by specialised cells in the gill, ionocytes, containing ion pumps and transporters ^57,58^. As a result, there are differential gill-specific transcriptional responses to salinity exposure in different tilapia species, triggering turnover of ionocytes as well as immune and cell stress response ^59^. We therefore suggest that the interactions of both STAT1-*prlr1* and SREBF2-*cldnh* are likely involved in salinity acclimation along the tilapia phylogeny. Discrete rewiring of existing ancestral variation at these important functional loci could be a mechanism to allow populations to rapidly adapt to new ecological niches e.g., *O. esculentus* introductions to Lake Rukwa ^60^, and also form the basis for breeding genotypes of desirable traits into farmed strains.

This novel epigenome of not only Nile tilapia gill, but the first recognised assessment of open chromatin in any fish gill tissue, will enable a better functional annotation of noncoding regions to identify causative variants associated with aquaculture relevant traits. Now that the ATAC-seq protocol has been optimised for tilapia fish, we can characterise the open chromatin landscape in the other 26 tilapia species to confirm and prioritise variants identified in this study, and define trait loci for other environmental traits e.g., temperature tolerance. The novel methods and resources generated here can be ultimately used to guide genomic selection programmes aimed at the genetic improvement of species to acclimate to adverse environmental conditions.

## Materials and Methods

### Tissue dissection

All animal procedures were approved by University of Stirling Animal Welfare and Ethical Review Body (AWERB) and carried out in accordance with approved guidelines. Three male *O. niloticus* individuals were sacrificed according to Home Office schedule 1 killing using overdose of MS-222 (tricaine) at University of Stirling, UK. The gill raker, arch and filaments were dissected from each individual, and one half was stored in RNAlater (1:5 ratio) for naked DNA and RNA extraction, and the other half immediately used for ATAC cell preparation.

### Cell preparation

Around 50,000 cells were harvested from the gill filaments by taking a 1mm biopsy punch, counting nuclei and then spun at 500×g for 5 min at 4°C. Cells were washed once with 50μl of ice-cold 1× PBS buffer and pelleted at 500×g for 5 min at 4°C. Cell pellets were gently resuspended in 50μl of 0.05% cold lysis buffer (10mM Tris-HCl pH7.4, 10mM NaCl, 3mM MgCl_2_, 0.05% IGEPAL CA-630), and immediately pelleted at 500×g for 10 min at 4°C and then stored on ice.

### Transposition reaction and purification

Pelleted nuclei were gently resuspended in a 50 μL transposition reaction mix composed of 25 μL 2× TD Buffer (Illumina), 2.5 μL Tn5 Transposes (Illumina) and 22.5 μL Nuclease-Free H_2_O. Transposition reaction was incubated at 37°C for 30 min and purified using the PCR purification MinElute Kit (QIAGEN), according to manufacturer’s protocol. Transposed DNA was eluted in 10 μL Elution Buffer (10mM Tris buffer, pH 8) and stored at -20°C.

### PCR amplification

Transposed DNA was amplified in a final volume of 50 μL composed of 10 μL Transposed DNA, 10 μL Nuclease Free H_2_O, 2.5 μL 25μM P7 adapter, 2.5 μL 25μM P5 adapter and 25 μL NEBNext High-Fidelity 2x PCR Master Mix (NEB). Transposed DNA was amplified for 5 min at 72°C, 30 secs at 98°C, and 11 cycles of 10 secs at 98°C, 30 secs at 63°C and 1 min at 72°C in a PCR thermocycler. Amplified transposed DNA was purified using the PCR purification MinElute Kit (QIAGEN) and eluted in 20 μL Elution Buffer (10mM Tris buffer, pH 8), according to manufacturer’s protocol. To remove excess primers for final ATAC libaries, an additional 1× Agencourt AMPure XP bead (Beckman Coulter) clean-up was performed and eluted in 0.1× filtered TE. ATAC libraries were quantified on the Qubit 4 fluorometer (Invitrogen) and size distribution assessed on Agilent Tapestation and/or Bioanalyser.

### DNA extraction

DNA was purified from gill filaments using the DNeasy Blood and Tissue kit (QIAGEN) according to manufacturer’s protocol. DNA was quantified on the Nanodrop 2000 (Thermo Scientific) and Qubit 4 fluorometer (Invitrogen), and used 1 ng for DNA library preparation using the Nextera XT DNA Library Preparation kit (Illumina), according to manufacturer’s protocol. Library size distribution was assessed on Agilent Tapestation and/or Bioanalyser. DNA libraries, obtained from naked DNA, were used as internal controls to determine background levels of genomic DNA accessibility and Tn5 transposase sequence cleavage bias.

### ATAC and control DNA sequencing

Three ATAC and three corresponding naked DNA control libraries were equimolar pooled and 50 bp paired-end sequenced at Earlham Institute on the Illumina NovaSeq 6000 platform using an S2 flow cell, generating an average of 90 million (ATAC-seq) and 16 million (naked DNA control) reads per library.

### ATAC-seq and control DNA processing

Sequence adaptors were removed and trimmed for quality from raw paired-end reads using Trim Galore! (v 0.6.5) (https://github.com/FelixKrueger/TrimGalore) and FastQC (v 0.11.9) ^61^ using default parameters. Read alignment, post alignment filtering and ATAC peak calling were performed according to the ENCODE projects ‘ATAC-seq Data Standards and Processing Pipeline’ for replicated data (https://www.encodeproject.org/atac-seq/). Briefly, trimmed reads were mapped to the Nile tilapia reference genome (*O. niloticus* UMD_NMBU) ^26^ using bowtie2 (v2.2.6) ^62^, with parameters `‘–k 4 –X2000 –mm`’ and outputted in BAM format using SAMtools (v1.9) ^63^. Since ATAC-seq generates high proportions (15-50% in a typical experiment) of mitochondrial mapped reads, any reads mapping to the mitochondrial genome were identified using BLAST (v2.3.0) ^64^ and removed from the BAM file using SAMtools (v1.9) ^63^. The resulting BAM files were sorted, and duplicated reads were marked using Sambamba v0.6.5 ^65^. Duplicated, unmapped, non-primary alignment, and failing platform QC reads were filtered out using SAMtools (v1.9) ^63^, retaining reads mapped as proper pairs, and fragment length distributions were plotted using Picard (v1.140) (https://github.com/broadinstitute/picard/). At each step, the recommended parameters from the ENCODE pipeline were applied. BAM files were converted to *tagalign* files using Bedtools (v2.30.0) ^66^ and Tn5 shifting of ATAC mappings carried out prior to peak calling. Peaks were identified using macs2 (v2.1.1) ^67,68^ with the shifted tag as test and corresponding control DNA as input with parameters `‘-f BED -p 0.05 --nomodel --shift -75 --extsize 150`’. Narrow peaks were used to create coverage tracks using *bedClip* and *bedToBigBed* in the UCSC-tools package (v333) (http://hgdownload.cse.ucsc.edu/admin/exe/). Following the ENCODE pipeline, Irreproducible Discovery Rate (IDR) peaks of true replicates were flagged as either true (<0.1) or false (≥0.1) using idr v2.0.4 (https://github.com/kundajelab/idr), taking reproducible (true) peaks between replicates. The fraction of reads in peaks (FRiP) were calculated using Bedtools (v2.30.0) ^66^ and demarcated as pass (>0.3) or acceptable (>0.2) according to ENCODE guidelines. FRiP was not used as a QC measure and instead, transcription start site (TSS) enrichment was calculated using ATACseqQC (v1.18.0) ^69^, with the TSS enrichment QC requirement of a significant signal value at the centre of the distribution being applied. All samples passed this criterion for further analysis. A union of peaks from all three biological replicates was created and used for subsequent analyses.

### Identifying conserved noncoding elements (CNEs)

We use a similar approach as applied previously ^5^ to call CNEs in Nile tilapia based on evolutionary constraint in a five cichlid species phylogeny. A multiple genome alignment (MGA) of four cichlid genomes (*M. zebra* UMD2a ^70^; *P. nyererei* v1 ^5^; *A. burtoni* v1 ^5^; *N. brichardi* v1 ^5^) and Nile tilapia (*O. niloticus* UMD_NMBU ^26^) was created using Cactus (v2.0.3) ^71^ and outputted in Multiple Alignment Format (MAF). A neutral substitution model was created using the *phyloFit* function of PHAST (v1.5) ^72^ by fitting a time reversible substitution ‘REV’ model and parameters ‘`—tree “((((Metriaclima_zebra,Pundamilia_nyererei),Astatotilapia_burtoni),N eolamprologus_brichardi),Oreochromis_niloticus)” --subst-mod REV’.` The five cichlid MGA was split by *O. niloticus* chromosomes/scaffolds using the *mafSplit* function in the UCSC-tools package (v333) (http://hgdownload.cse.ucsc.edu/admin/exe/). The neutral substitution model and MGAs of each *O. niloticus* reference chromosome/scaffold were used as input to predict conserved noncoding elements (CNEs) using the *phastCons* function of PHAST (v1.5) ^72^ with parameters `‘--target-coverage 0.3 --expected-length 30 --most-conserved -- estimate-trees --msa-format MAF`’. Conservation score outputs were used to define highly conserved CNEs (hCNEs) with sequence identity ≥90% over ≥30bp and pseudo accelerate/diverged CNEs (pseudo aCNEs) with sequence identity <90% over ≥30bp. The evolutionary conservation-acceleration (CONACC) score of each pseudo aCNE was predicted by running *phyloP* ^73^ of the PHAST (v1.5) package ^72^ using the likelihood ratio test (LRT) ‘`--method LRT`’ on the CNE `-- features` with their corresponding neutral substitution model in ‘`--mode CONACC`’. Pseudo aCNEs with a negative CONACC score, likelihood ratio <0.05, and significant divergence (altsubscale >1), were defined as significantly deviating from the neutral model, and therefore as true aCNEs.

### Peak annotation and enrichment

Up to 5 kb gene promoter regions were annotated in the *O. niloticus* UMD1 ^26^ genome according to the method used in our previous study ^7^. Narrow peaks either overlapping or most proximal to all annotated features ^26^ in the genome were mapped using the *intersect* function of Bedtools (v2.30.0) ^66^, and each peak assigned to a feature and/or gene accordingly. The enrichment of accessible peaks in coding and noncoding regions was tested using the Genome Association Tester (GAT) tool ^74^. The accessible peaks were provided as segments of interest to test, a bed file of annotations to test against, and workspace as the length of each scaffold/chromosome. GAT was ran using the following parameters: `--verbose=5, --counter=segment-overlap, --ignore-segment-tracks, --qvalue-method=BH --pvalue-method=norm`. We use the Benjamini-Hochberg ^75^ false discovery rate (FDR) to assess enrichment of peaks in annotated regions, with a statistical cut-off of FDR < 0.05.

### Gene Ontology (GO) enrichment

GO enrichment analyses of genes was conducted using the ‘g:GOst’ module of g:Profiler (https://biit.cs.ut.ee/gprofiler/gost) ^76^, version e105_eg52_p16_e84549f (February 2022), using the *O. niloticus* database. We use the false discovery rate (FDR) corrected hypergeometric *p*-value to assess enrichment of GO terms, with a statistical cut-off of FDR < 0.05.

### Transcription factor (TF) footprinting

TF footprints were characterised using HINT-ATAC in the Regulatory Genomic Toolbox (v0.13.0) ^77^ using a stringent false positive rate (FPR) of 0.0001, with both Nile tilapia specific and cichlid-wide position weight matrices (PWMs) as defined in our previous study ^7^, as well as vertebrate PWMs from JASPAR (v9.0) ^78^, HOCOMOCO ^79^, GTRD ^80^, and UniPROBE ^81^. TF footprint line plots were generated with the ‘differential analysis’ module of HINT-ATAC in the Regulatory Genomic Toolbox (v0.13.0) package ^77^ using bias-corrected signals. Redundant TFBSs in the same, or overlapping positions were filtered based on selecting the highest bit-score of motif match for each overlapping TF.

### RNA extraction and sequencing

RNA was purified from each tissue using the RNeasy Plus Mini kit (QIAGEN) according to manufacturer’s protocol. RNA and DNA content were quantified on the Qubit 4 fluorometer (Invitrogen) and integrity assessed on Agilent Tapestation and/or Bioanalyser, taking samples with RIN≥7 and <15% genomic DNA. A total of 45/55 (82%) samples passed these criteria for selection (Supplementary Table S2). A total of 45 stranded RNA libraries were prepared using the NEBNext Ultra II Direction RNA-seq kit according to manufacturer’s protocol. All stranded RNA-seq libraries were equimolar pooled and 150 bp paired-end sequenced at Earlham Institute on 1 lane of the Illumina NovaSeq 6000 platform using an S4 v1.5 flow cell, generating an average of 70 million reads per library.

### RNA-seq processing

Read quality was assessed using FastQC (v 0.11.9) ^61^ and Trim Galore! (v 0.6.5) (https://github.com/FelixKrueger/TrimGalore) was used to remove adapters and trim for low-quality from the raw paired-end reads using default settings. All reads were then mapped to the *O. niloticus* UMD1 genome ^26^ using HISAT2 (v 2.2.1) ^82^ with default parameters. Mapping QC was carried out using QualiMap (v 2.2.1) ^83^ with default ‘rnaseq’ parameters. The final BAM file was sorted using samtools (v1.16.1) ^63^ and transcript abundance was calculated using ‘htseq-count’ in the HTSeq (v 2.0.2) ^84^ package.

### Identifying ATAC peak and gene expression association

Given the large number of accessible peaks and overlapping genes (127,602 peaks and 20,248 genes), we devised an approach to reduce and identify the peaks that could regulate their gene targets expression. For gene expression, we use an approach applied previously ^85^ to calculate transcript per million (TPM) values by first normalizing the transcript count by the gene length, as calculated using GTFtools ^86^, followed by the library size, as calculated using QualiMap (v 2.2.1) ^83^, and then carrying out a *log2*(x +1) transformation of 1) each genes TPM in each replicate; and 2) the mean TPM for each gene across biological replicates. Each genes mean TPM across the three replicates is used to assess correlations with ATAC signal whereas replicate specific TPM are used to study any differences between replicates. The ATAC signal was processed using an approach applied previously ^87^ to obtain the number of independent Tn5 insertions in collated gene promoter peaks; the number of insertion sites (peak counts) was counted using collated narrow peaks against a merged BAM file of all three replicates using Bedtools (v2.30.0) *coverage* ^66^. After creating a peak count matrix, the counts matrix was normalised using edgeR ^88^ counts per million (CPM) ‘`log=TRUE, prior.count=5`’ followed by a quantile normalization using the preprocessCore (https://github.com/bmbolstad/preprocessCore) *normalize*.*quantiles* module in R (v 4.4.2). After merging the three replicates ATAC signal using the *log2* average from the normalized counts matrices, the average *log2*-TPM and *log2*-ATACSignal for each gene was plotted with the correlation co-efficient (r) calculated in R (v 4.4.2).

Based on an approach devised previously ^30^ to categorise putative activated, repressed or poised genes, peak-gene relationships are assigned to one of four groups according to lower than 50th (medium-low) and higher than 70th (high) percentiles of gene promoter accessibility (*log2*-ATACSignal) and gene expression (*log2*-TPM). The four groups are (1) MA–ME (medium–low accessibility and medium–low expression); (2) HA–HE (high accessibility and high expression); (3) HA–ME (high accessibility and medium–low expression); (4) MA–HE (medium–low accessibility and high expression). Any genes that did not fall into one of the four groups were not used in the analysis to maintain stringent gene groups.

### Identifying SNPs with exclusive variant types between Nile tilapia and other species

The VCF file of the 69 million SNPs mapped to the Nile tilapia (*O. niloticus* UMD_NMBU ^26^) reference genome in our previous study (Ciezarek, A. *et al*., bioRxiv TBC, 2023) was first normalised to split multiallelic sites into multiple rows using bcftools (v 1.12) ^63^ with the parameters ‘`norm -m-any -Ov`’. After outputting the first four columns from the resulting file, we use bcftools ‘`query`’ to tabulate the number of samples having each variant type, and then bcftools ‘`view`’ to report each sample with each type of variant. SNPs that are present in prioritised TFBSs of candidate genes are then filtered for variant types (heterozygous and/or homozygous alternate) that are exclusive to Nile tilapia and any other species.

## Declarations

### Data Availability

All sequencing data generated for this article is available under the ENA study accession PRJEB59919. All other data underlying this article is available in the article, its supplementary material or in the following repository: https://doi.org/10.6084/m9.figshare.22100825.

## Acknowledgments

The authors acknowledge the support of the Biotechnology and Biological Sciences Research Council (BBSRC), part of UK Research and Innovation. The authors would like to acknowledge Tom Barker, Vanda Knitlhoffer, Leah Catchpole, and Suzanne Henderson of the Genomics Pipelines Group at Earlham Institute for data generation including preparation of RNA-Seq libraries, pooling, and sequencing. The authors would also like to acknowledge the Scientific Computing group, as well as support for the physical HPC infrastructure and data centre delivered via the NBI Research Computing group.

## Author contributions

KR and DP sacrificed and dissected fresh fish tissues; TKM prepared tissue-specific cells to perform transposition reactions and purifications for ATAC libraries; TKM and AM performed tagmentation of genomic DNA controls, library amplification and QC; AC identified variants in the tilapia phylogeny; TKM processed ATAC-seq and RNA-seq data, identified CNEs, annotated peaks with enrichment analysis, ran GO enrichment, identified TF footprints, and identified ATAC peak and gene expression associations; TKM and WH wrote the manuscript with input from AM, AC, KR, DP and FDP.

