## Supplementary Information for "Chromatin accessibility associated with aquaculture relevant traits in tilapia"

### Results

#### Accessible gene promoter regions in gill tissue are functionally associated with aquaculture relevant traits

Despite having the smallest enrichment of accessible peaks (Fig. 1), we focus on gene promoter regions, taken as up to 5 kb from the transcription start site (TSS) [1], because they 1) have a similar enrichment of SNPs in accessible peaks in intronic regions (both with a fold enrichment of 1.1, adjusted *P*-value <0.05), and higher compared to all other noncoding regions (fold enrichment of 0.8 – 1.08, adjusted *P*-value <0.05); 2) harbour regulatory binding sites e.g., TFBSs that have functionally diverged [1,2], and can be associated with gene expression differences [1] in Nile tilapia and East African cichlids; and 3) can be associated to their neighbouring gene for functional analyses.

Out of the 301,293 open chromatin (accessible) peaks in gill tissue, 127,602 accessible peaks (average size of 464 bp) were found within gene promoters of 69% (20,248 genes) of all 29,552 [3] annotated Nile tilapia genes. The 9,304 missing genes are enriched (false discovery rate [FDR] < 0.05) for several immune system and response related gene ontology (GO) biological processes (Supplementary Fig. S1 – S2), that could be better associated to lymphoid tissues e.g., kidney and spleen that are not sampled in this study.

#### Nile tilapia SNPs are enriched in noncoding regions

20,101 (99%) of all 20,248 accessible gene promoters have SNPs in accessible regions. Most SNPs are rare variants (Minor Allele Frequency, MAF <= 0.1) with the largest percentage of rare variants found in gene promoter (4,924,352/5,470,066 – 90%) and accessible gene promoter (980,565/1,083,195 – 91%) regions compared to the whole genome (59,995,966/69,064,774 – 87%) (Supplementary Fig. S4a-b). We find that 1% more rare variants (Minor Allele Frequency, MAF <= 0.1) are found in noncoding regions that are accessible (88% MAF<= 0.1 and 70% MAF <=0.03) than non-accessible (87% MAF<= 0.1 and 68% MAF <=0.03) (Supplementary Fig. S4c-d).

**Discrete variation in accessible TFBSs is likely driving gene expression associated with tilapia gill adaptations**

We previously focused on TFBSs between Nile tilapia and four other cichlids [1,2] however, these were *in silico*-based predictions using the genome sequence, and not chromatin accessibility. By focusing on TFBSs (derived from footprints, see '*Materials and Methods*') of accessible gene promoter regions, we first characterised transcription factor (TF) – target gene (TG) relationships. In the Nile tilapia gill, we identified 3,779,403 TF footprints and using 2042 TF motifs, we predicted a total of 40,007,918 TFBSs (including overlapping sites of the same motif) across the whole genome (see '*Materials and Methods*'). When filtering for redundant motifs in overlapping positions (see '*Materials and Methods*'), we identified a total of 23,955,801 (60% of the 40,007,918) predicted TFBSs across the whole genome.

### Supplementary Figures

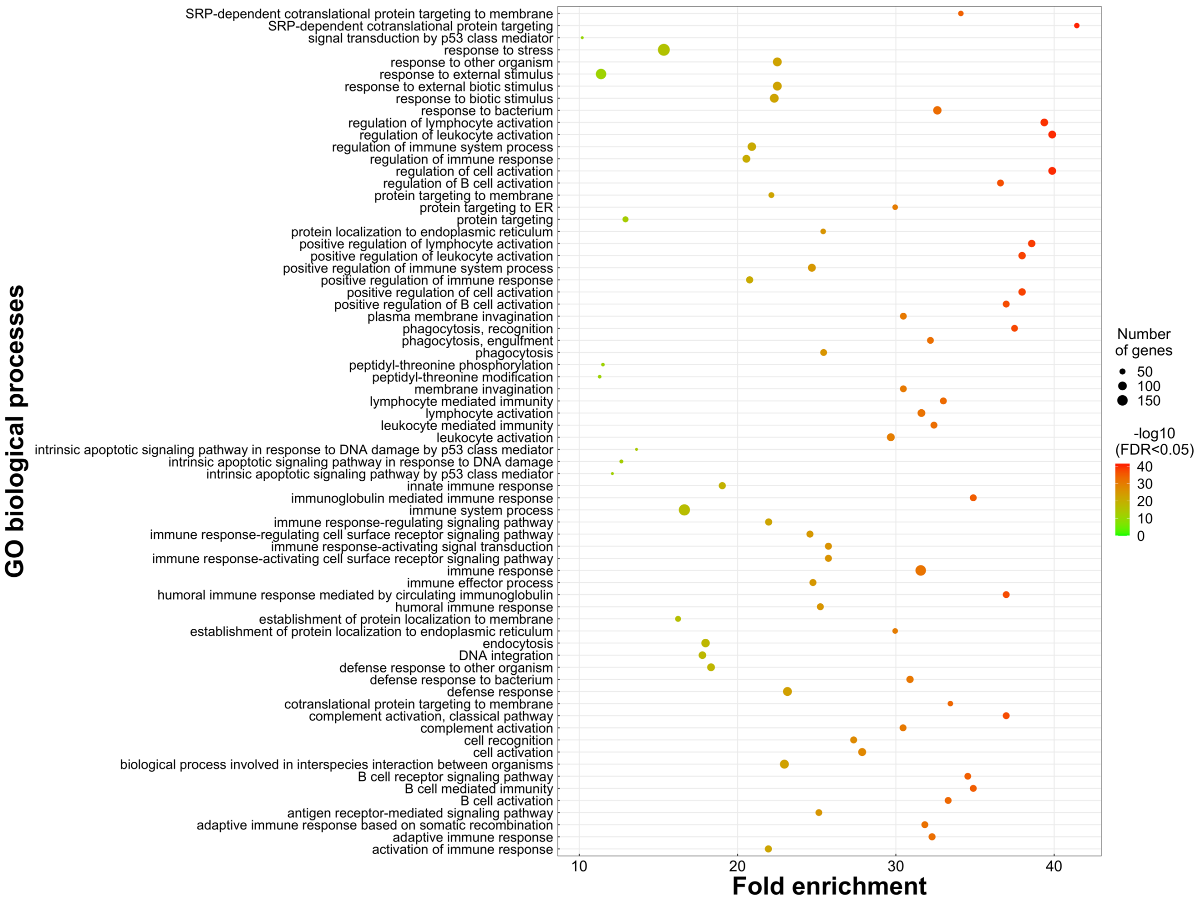


**Supplementary Fig. S1 – Topmost enriched ontology terms of 9,304 genes without an accessible peak in Nile tilapia gill tissue .** GO terms are subset for enrichment of -log10 (FDR < 0.05) of more than 10, with the full figure provided as Supplementary Fig. S2. Circles show enriched term (y-axis) of significance (FDR <0.05, heatmap to right) and fold enrichment (x-axis) values of GO terms. Number of genes missing an accessible peak are shown by the size of each circle.

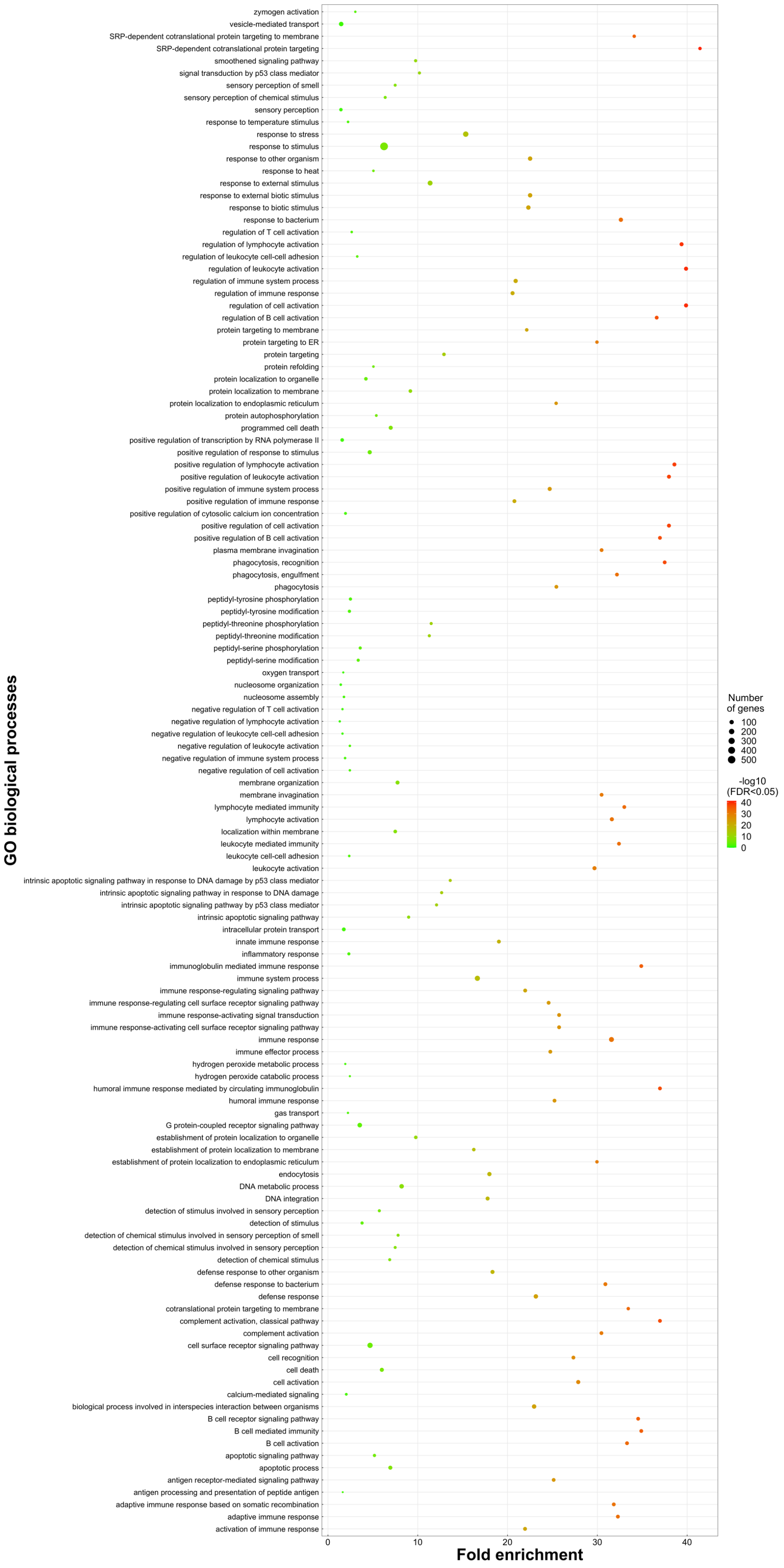


**Supplementary Fig. S2 – Enriched ontology terms of 9,304 genes without an accessible peak in Nile tilapia gill tissue .** Circles show enriched term (y-axis) of significance (FDR <0.05, heatmap to right) and fold enrichment (x-axis) values of GO terms. Number of genes missing an accessible peak are shown by the size of each circle.

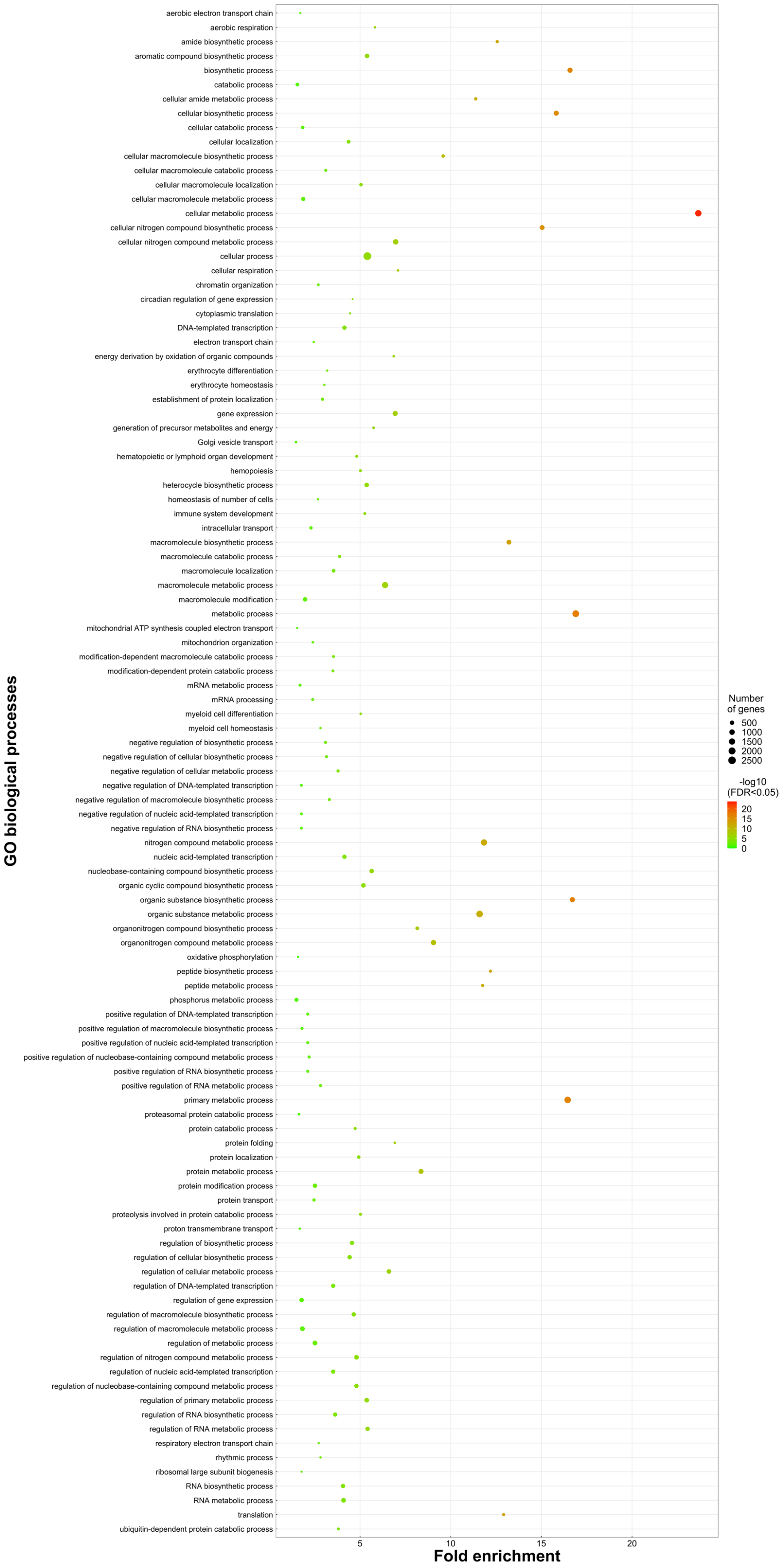


**Supplementary Fig. S3 – Enriched ontology terms of 4,117 genes with high accessibility and high expression (HA-HE) in Nile tilapia gill tissue .** Circles show enriched term (y-axis) of significance (FDR <0.05, heatmap to right) and fold enrichment (x-axis) values of GO terms. Number of genes missing an accessible peak are shown by the size of each circle.

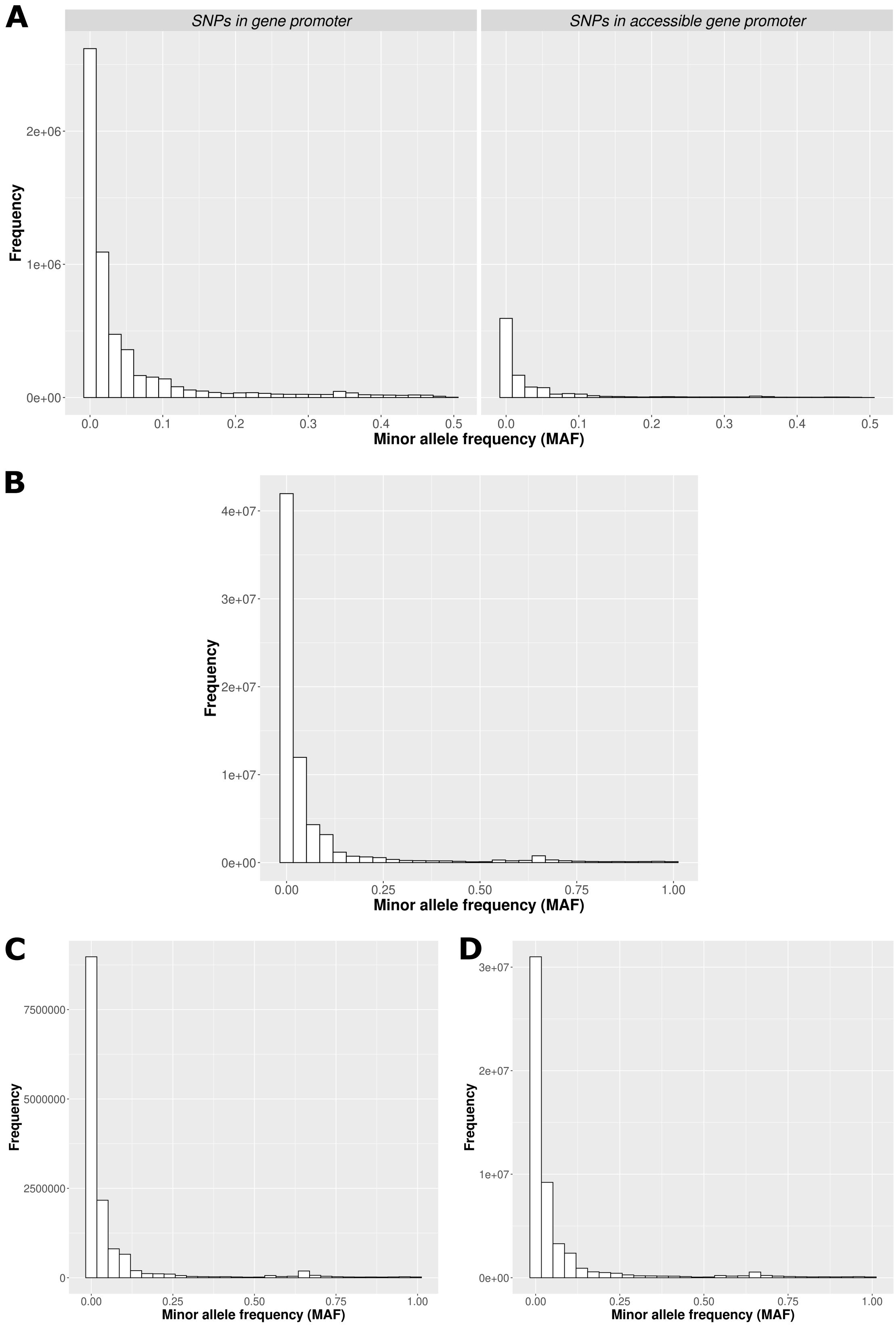


**Supplementary Fig. S4 – Frequency of Minor Allele Frequencies (MAF) of SNPs in different regions of the Nile tilapia genome.** Histograms show the frequency (y-axis) and MAF of **(A)** SNPs in all up to 5 kb gene promoter regions (left) and accessible gene promoter regions that have reproducible peaks between all three gill replicates (right). **(B)** genome-wide SNPs. **(C)** noncoding SNPs in accessible (open chromatin) regions. **(D)** noncoding SNPs in non-accessible regions.

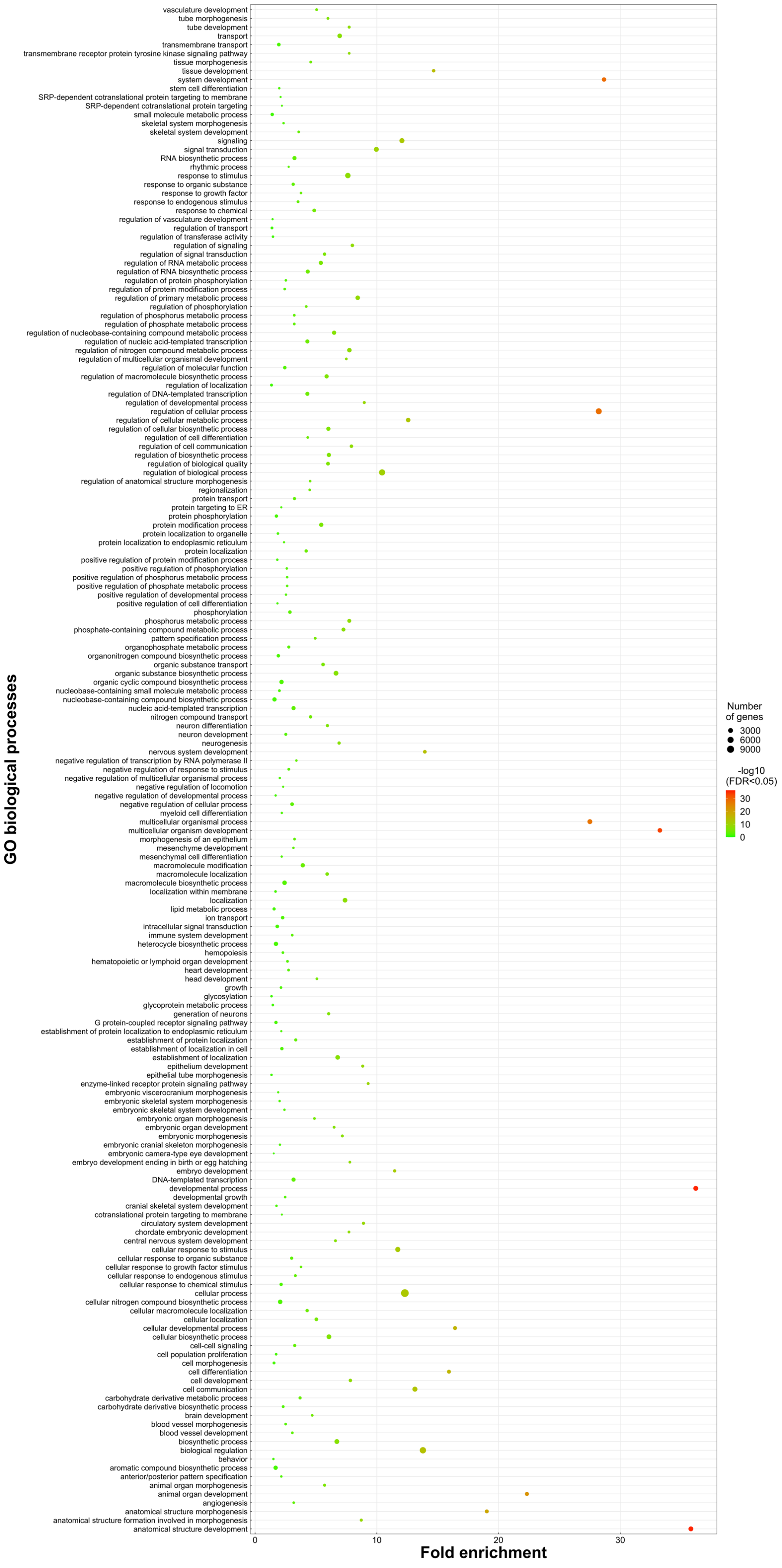


**Fig. S5 – Enriched ontology terms of 20,248 accessible gene promoter regions containing SNPs in the Nile tilapia genome.** Circles show enriched term (y-axis) of significance (FDR <0.05, heatmap to right) and fold enrichment (x-axis) values of GO terms. Number of genes with promoter regions containing SNPs shown by size of each circle.
